# Sulcal depth in prefrontal cortex: A novel predictor of working memory performance

**DOI:** 10.1101/2021.11.03.467157

**Authors:** Jewelia K. Yao, Willa I. Voorhies, Jacob A. Miller, Silvia A. Bunge, Kevin S. Weiner

**Author notes:** Corresponding authors: Kevin S. Weiner 2121 Berkeley Way West Berkeley, CA USA 94720, Silvia A. Bunge 2121 Berkeley Way West Berkeley, CA USA 94720.

## Abstract

The neuroanatomical changes that underpin cognitive development is of major interest in neuroscience. Of the many aspects of neuroanatomy to consider, tertiary sulci are particularly appealing as they emerge last in gestation, show a protracted development after birth, and are either human- or hominoid-specific. Thus, they are ideal targets for exploring morphological-cognitive relationships with cognitive skills, such as verbal working memory (WM), that also show protracted development. Yet, the relationship between sulcal morphology and verbal WM is unknown, either in development or more generally. To fill this gap, we adopted a data-driven approach with cross-validation to examine the relationship between sulcal depth in lateral prefrontal cortex (LPFC) and verbal WM in 60 participants ages 6-18. These analyses identified nine left, but not right, LPFC sulci (of which six were tertiary) whose depth predicted verbal WM performance. Most of these sulci are located within and around contours of functionally defined parcellations of LPFC proposed previously. This sulcal depth model out-performed models with age or cortical thickness. Taken together, these findings contribute to building empirical support for a classic theory that tertiary sulci serve as landmarks in association cortices that contribute to aspects of human behavior that show a protracted development.

## **1.** Introduction

A fundamental point of exploration in developmental cognitive neuroscience is the relationship between brain anatomy and cognition. For instance, many studies have examined how individual differences in cognitive development relate to cortical thickness, volume, and/or changes in white matter tissue properties (e.g., Bathelt et al. 2018; Darki and Klingberg, 2015; Dickerson et al., 2008; Lu et al., 2007; Øtsby et al., 2011; Shaw et al., 2006; Tamnes et al., 2013; Wendelken et al., 2017; Yeatman et al., 2012). To a much lesser extent, previous studies have also examined the relationship between cognition and sulcal morphology in children (Cachia et al., 2018; Chung et al., 2017; Gregory et al., 2016; Tissier et al., 2018). Importantly, little is known about the role in cognitive development (or cognition more broadly) of shallow, variable cortical indentations known as tertiary sulci, which often must be individually identified (Amiez et al., 2018; Paus et al., 1996; Petrides, 2019).

Tertiary sulci are particularly interesting for three main reasons. First, they emerge late in gestation and continue to develop after birth (Petrides, 2019; Sanides, 1962, 1964; Weiner, 2019; Welker, 1990). Second, they are largely hominoid-specific structures and are particularly prominent in human association cortices such as lateral prefrontal cortex (LPFC), which supports higher-level cognition (Duncan and Owen, 2000; Passingham and Wise, 2012; Stuss and Knight, 2002). Third, due to the protracted development of both tertiary sulci and higher order cognitive skills associated with LPFC, Sanides (1962, 1964) proposed a classic hypothesis that tertiary sulci likely serve as anatomical and functional landmarks that are behaviorally relevant in association cortices such as LPFC. In recent years, there has been mounting evidence in support of Sanides’ hypothesis in ventral temporal cortex across the lifespan (VTC; Weiner, 2019), as well as in medial PFC (Lopez-Persem et al., 2019) and in LPFC in both adults (Miller et al., 2021) and children (Voorhies et al., 2021). Directly relevant for the present study, the latter findings— supported by the notion that depth is a characteristic feature of tertiary sulci, which are much more shallow than primary and secondary sulci (Armstrong et al., 1995; Miller et al., 2021; Weiner, 2019)—showed that the depth of a subset of right-lateralized LPFC tertiary sulci correlated with performance on a visuospatial reasoning task.

Here, we build on those findings to test for a relationship between the depth of LPFC tertiary sulci and verbal working memory (WM) – a higher-level cognitive ability that depends in part on LPFC (e.g., Fuster, 2001). WM *maintenance* refers to the ability to keep mental representations active over the short-term: for example, rehearsing a phone number in verbal (or phonological) WM (Baddeley and Hitch, 1974). By contrast, WM *manipulation* refers to the ability to reorganize or transform - i.e., ‘work with’ - these active representations (Goldman-Rakic, 1995). WM ability, particularly manipulation, improves over childhood and adolescence (Cowan, 2016; Gathercole et al., 2004), and this improvement has been linked to LPFC development (Goldman-Rakic, 1987; for review, see Bunge and Wright, 2007).

Although neuroscientific research has broadly implicated LPFC in WM, LPFC is a large and highly heterogeneous area. Brain imaging studies have reported functional dissociations along both the dorsal-ventral and rostral-caudal axes (Petrides, 2005; Blumenfeld et al., 2013; Badre,2008; D’Esposito et al., 1999; Fuster, 2004; Koechlin, Ody, and Kouneiher, 2003; Owen et al., 1996). With respect to verbal WM, specifically, neuroimaging studies have drawn a distinction between the inferior frontal gyrus (IFG), also known as ventrolateral PFC (VLPFC; Brodmann areas 44, 45, 47), and the middle frontal gyrus (MFG) - specifically, a portion known as mid-dorsolateral PFC (mid-DLPFC; Brodmann areas 9/46). Left IFG has been strongly linked to verbal, or phonological, WM capacity (Amiez and Petrides, 2007). By contrast, bilateral MFG has been implicated in control processes that operate on the contents of WM - that is, WM manipulation (D’Esposito et al., 1999; Owen et al., 1996; Sakai and Passingham, 2003; Crone et al., 2006) and monitoring (Amiez and Petrides, 2007). However, our understanding of the role of distinct LPFC subregions in verbal WM is limited by the fact that PFC lesions in humans tend to be quite large, and because insufficient attention is paid to sulcal anatomy in the WM neuroimaging literature (but see Amiez and Petrides, 2007) - let alone to sulcal variation among individuals.

The present study examines the relationship between sulcal anatomy in LPFC and verbal WM in children and adolescents. Based on our previous findings with regards to visuospatial reasoning (Voorhies et al., 2021), we hypothesized that the depth of a subset of LPFC sulci would be linked to verbal WM performance during development. We had three predictions. First, given Sanides’ hypothesis, we predicted that the depth of the shallow, late-developing tertiary sulci would be particularly predictive of individual differences in verbal WM. Second, given ample evidence that verbal WM is left-hemisphere dominant, we predicted a leftward asymmetry in sulcal-cognitive relations. Third, given previously documented dorsal-ventral dissociations within LPFC with regards to verbal WM, we predicted that sulci in the IFG/VLPFC would be broadly implicated in verbal WM maintenance and manipulation, whereas those in the MFG/mid-DLPFC would be particularly linked to WM manipulation.

To directly address our predictions, our study aimed to answer three main questions: 1) Is there a relationship between verbal WM maintenance and/or manipulation and mean depth of LPFC sulci? 2) If so, do these relationships differ as a function of type of sulcus (shallow/tertiary or deep/non-tertiary), hemisphere, and/or task demands (WM maintenance and/or manipulation)? 3) Can we construct a model to predict an individual’s verbal WM task score from sulcal depth? To answer these questions, we manually defined 2,157 sulci in 60 participants between ages 6 and 18. Altogether, the present study establishes a novel link between LPFC sulcal morphology and verbal WM skills in a pediatric cohort.

## 2. Materials and Methods

### 2.1. Participants

Our present analyses leverage previously published data from the Neurodevelopment of Reasoning Ability (NORA) study (e.g., Wendelken et al., 2016; 2017). 60 typically developing children and adolescents (all right-handed native English speakers; 27 identified as female, 33 identified as male) were randomly selected from the dataset (see Supplementary Fig. 2 for demographic information). 48 of these participants were also included in Voorhies et al. (2021). Participants ranged in age from 6 to 18 (*M* = 12.12, *SD* = 3.39; Females: *M* = 11.82, *SD* = 3.23; Males: *M* = 12.36, *SD* = 3.50). All participants were screened for neurological impairments, psychiatric illness, history of learning disabilities, and developmental delay. All participants and their parents gave informed assent or consent to the study, which was approved by the Committee for Protection of Human Subjects at the University of California, Berkeley.

### 2.2. Data Acquisition

#### 2.2.1. Behavioral Data

Verbal WM was measured via raw scores of the Digit Span task from the 4th edition of the Wechsler Intelligence Scale for Children (WISC-IV; Wechsler, 1974). The Digits Forward condition of the Digit Span task taxes WM maintenance, whereas the Backward condition taxes both WM maintenance and manipulation. In Digits Forward, the experimenter reads aloud a sequence of single-digit numbers, and the participant is asked to immediately repeat the numbers in the same order; in Digits Backward, they are asked to immediately repeat the numbers in the reverse order. The length of the string of numbers increases after every two trials. The Forwards task has eight levels, progressing from 2 to 9 digits (16 total trials). The Backwards task has seven levels, from 2 to 8 digits (14 total trials). Participants are given a score of 1 for a correct answer or a 0 for an incorrect answer. Testing on a given task continues until a participant responds incorrectly to both trials at a given level, after which the experimenter recorded a score out of 16 for Digits Forward and a score out of 14 for Digits Backward.

#### 2.2.2. Imaging Data Acquisition

Brain imaging data were collected at the UC Berkeley Brain Imaging Center on a Siemens 3T Trio system. High-resolution T1-weighted MPRAGE anatomical scans (TR=2300ms, TE=2.98ms, 1×1×1mm voxels) were acquired for cortical morphometric analyses.

#### 2.2.3. Cortical Surface Reconstruction

All T1-weighted images were visually inspected for scanner artifacts. Using FreeSurfer’s automated segmentation tools (FreeSurfer 6.0.0: http://surfer.nmr.mgh.harvard.edu; Dale et al., 1999), each anatomical volume was segmented to separate gray from white matter, and the resulting boundary was used to reconstruct the cortical surface for each subject. Each reconstruction was visually inspected for segmentation errors and manually corrected when necessary.

### 2.3. Morphological Analyses

#### 2.3.1. Sulcal labeling

Sulcal labels in LPFC were based on the most recent parcellation proposed by Petrides (2019; see also Sprung-Much and Petrides, 2018, 2020). In each hemisphere, LPFC sulci in the IFG/ VLPFC and MFG/mid-DLPFC were manually defined at the level of individual participants on both the pial and inflated surfaces (Fig. 1, Table 1), along with the *inferior frontal sulcus (ifs)*, which divides these two gyri. As controls, we also labeled two precentral sulcal components and the central sulcus, which have not been linked to WM.

**Figure 1.**
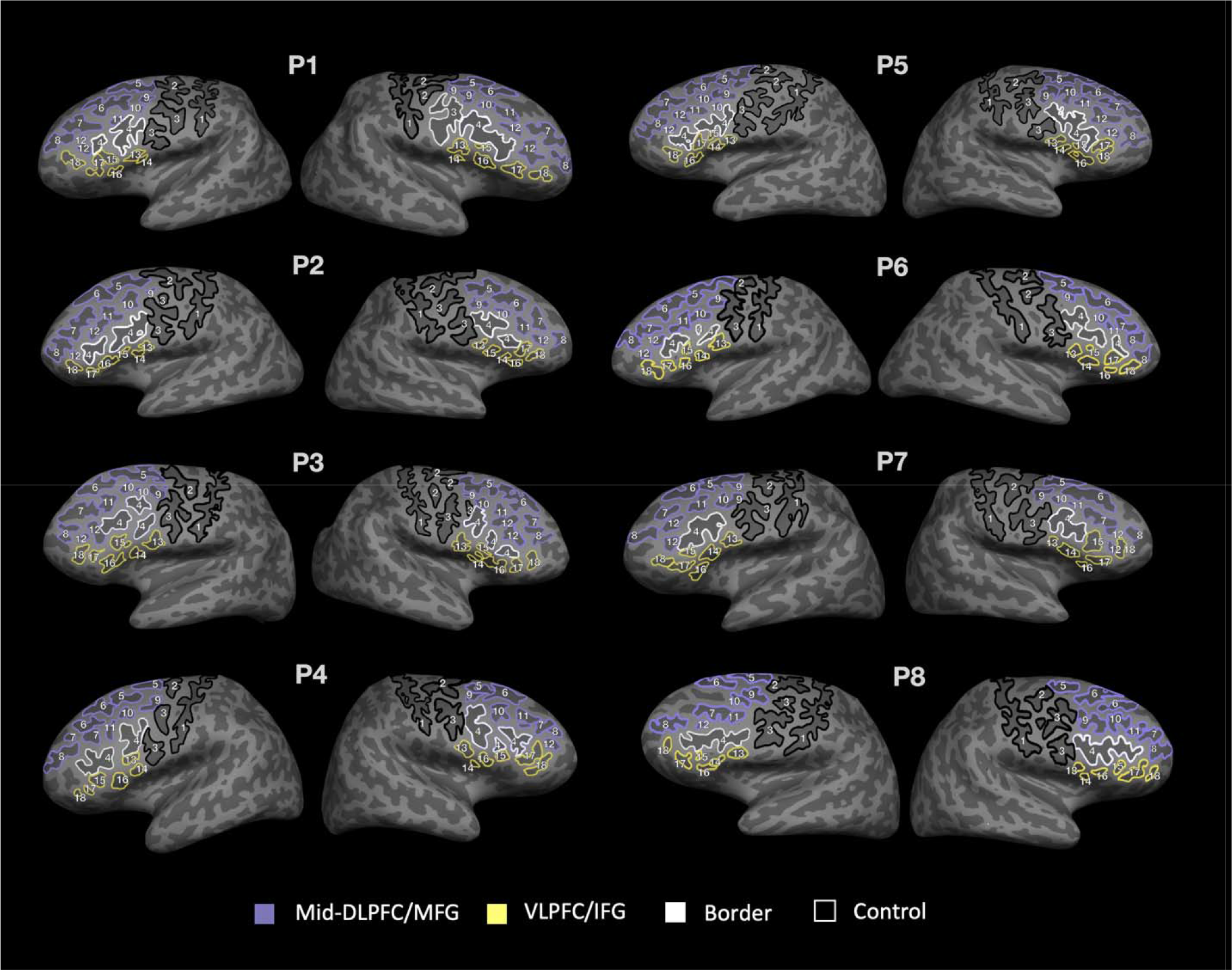
Sulcal definitions in lateral prefrontal cortex (LPFC) in 6-18 year-olds. Inflated surfaces of the left and right hemispheres in eight example participants (randomly chosen). 18 manually labeled sulci are outlined and labeled by number on each surface based on definitions derived from Petrides (2019). Eight sulci are included in MFG/mid-DLPFC (purple) and six in IFG/VLPFC (yellow). The ifs (white) is considered the boundary between mid-DLPFC and VLPFC. Sulci outlined in black (cs, sprs, iprs) are not included in these verbal WM regions and are considered as control sulci. The pimfs can be comprised of both a dorsal and a ventral component (RH: P1, P3, P5, P7; LH: P1, P2, P3, P5, P6, P7), include only one of these components (RH: P2, P4; LH: P8), or be absent (RH: P6, P8; LH: P4). See Supplementary Figure 1 for all 2,157 sulcal definitions in all participants. See Supplementary Figure 2 for demographic information for all participants.

**Table 1.** Sulcal definitions in LPFC. Table of sulci defined and outlined in each participant. Numbers correspond to those included in *Figure 1*. Sulcal abbreviations, full name, and verbal WM anatomical region are also included. Sulci not included in either the MFG/mid-DLPFC or IFG/VLPFC are defined as control (cs, iprs, sprs) or border (*ifs*) sulci. *The pimfs can have 0, 1, or 2 components; dorsal and ventral components have been combined into one label for the present study.

| No. | Abbreviation | Name | Classification |
| --- | --- | --- | --- |
| 1 | cs | central sulcus | Control |
| 2 | sprs | superior precentral sulcus | Control |
| 3 | iprs | inferior precentral sulcus | Control |
| 4 | ifs | inferior frontal sulcus | Border |
| 5 | sfs-p | superior frontal sulcus – posterior | MFG/mid-DLPFC |
| 6 | sfs-a | superior frontal sulcus – anterior | MFG/mid-DLPFC |
| 7 | imfs-h | intermediate frontal sulcus – horizontal | MFG/mid-DLPFC |
| 8 | imfs-v | intermediate frontal sulcus – vertical | MFG/mid-DLPFC |
| 9 | pmfs-p | posterior middle frontal sulcus – posterior | MFG/mid-DLPFC |
| 10 | pmfs-i | posterior middle frontal sulcus – intermediate | MFG/mid-DLPFC |
| 11 | pmfs-a | posterior middle frontal sulcus – anterior | MFG/mid-DLPFC |
| 12 | pimfs* | paraintermediate frontal sulcus | MFG/mid-DLPFC |
| 13 | ds | diagonal sulcus | IFG/VLPFC |
| 14 | aalf | ascending ramus of the lateral fissure | IFG/VLPFC |
| 15 | ts | triangular sulcus | IFG/VLPFC |
| 16 | half | horizontal ramus of the lateral fissure | IFG/VLPFC |
| 17 | prts | pretriangular sulcus | IFG/VLPFC |
| 18 | lfms | lateral frontomarginal sulcus | IFG/VLPFC |

The location and definition of each sulcus was confirmed by two trained independent raters (authors JKY and WIV) and then finalized by a neuroanatomist (KSW). The surface vertices for each sulcus were selected using tools in FreeSurfer and saved as surface labels for vertex-level analysis of morphological statistics. As it can sometimes be difficult to determine the precise start and end points of a sulcus on one surface (Borne et al., 2020), all definitions were also guided by the pial and smoothwm surfaces of each individual. Using multiple surfaces allowed us to form a consensus across surfaces and clearly determine each sulcal boundary as in our previous work (Miller et al., 2021; Voorhies et al., 2021).

The dorsal and ventral components of the *pimfs* were not identifiable in all participants. However, we could identify at least one component in the right hemisphere of 58/60 participants and in the left hemisphere of 59/60 participants. For our analyses, our inclusion criteria were to include participants who had at least one *pimfs* component (N=57). For participants who had both the dorsal and ventral components, we merged these components into one *pimfs* label using the FreeSurfer function *mris_mergelabels*. Thus, our findings are reported for the merged label. This process resulted in 2,157 manually defined sulci.

#### 2.3.2. Distinction among Sulcal Types

As described in our previous work, as well as classic studies, tertiary sulci are defined as the last sulci to emerge in gestation after the larger and deeper primary and secondary sulci (Bailey and Bonin, 1951; Bailey et al., 1950; Chi, 1977; Connolly, 1940, 1950; Cunningham, 1892; Miller, 2021a,b; Petrides, 2019; Retzius, 1896; Sanides, 1964; Turner, 1948; Weiner, 2014, 2016, 2019; Welker, 1990). Previous studies define primary sulci as sulci that emerge before 32 weeks in gestation, secondary sulci as those emerging between 32-36 weeks in gestation, and tertiary sulci as sulci that emerge during and after 36 weeks in gestation (Chi et al., 1977; Connolly, 1940, 1950; Cunningham, 1982; Miller et al., 2021a, 2021b; Retzius, 1896; Turner, 1948). Based on these definitions, we consider the *cs, sprs, iprs, sfs-p, sfs-a,* and *ifs* as primary sulci; and the *pmfs-p, pmfs-i, pmfs-a, pimfs, ds, ts, aalf, half, lfms,* and *prts* as putative tertiary sulci. The definitions of these latter ten sulci as putative tertiary sulci are further supported by their shallow depth (Figure 2), which is a defining morphological feature of tertiary sulci. Apart from these sulci, the question of whether or not other LPFC sulci should be considered secondary or tertiary is still unresolved. For example, the *imfs-v* and *imfs-h* are contemporary labels for classic definitions of sulci commonly labeled as either the *frontomarginal* and/or *middle frontal sulci* (Miller et al., 2021a,b; Petrides, 2019). When considering classic papers and atlases (Connolly, 1940; Cuningham, 1892; Retzius, 1896; Turner, 1948), both the *imfs-h* and *imfs-v* appear to be prevalent prior to 32 weeks, which would define them as primary sulci. Yet, additional studies define sulci in this cortical expanse as secondary (Tamraz and Comair, 2006). As in our previous studies (Miller et al., 2021a,b; Voorhies et al., 2021), we consider the *imfs-h* and *imfs-v* as primary sulci in the present study, but it is possible that future studies will establish them as secondary sulci. Of course, the definition of primary, secondary, and tertiary sulci remains contentious, which is why we refer to LPFC tertiary sulci explored in the present study as “putative.” Future research leveraging non-invasive fetal imaging will be sure to further improve the distinctions among primary, secondary, and tertiary sulci. Critically, our data-driven approach—and in turn, our findings—are agnostic to these distinctions. That is, the model-based approach adopted here quantitatively determines which sulci best predict working memory performance, regardless of their classification.

**Figure 2.**
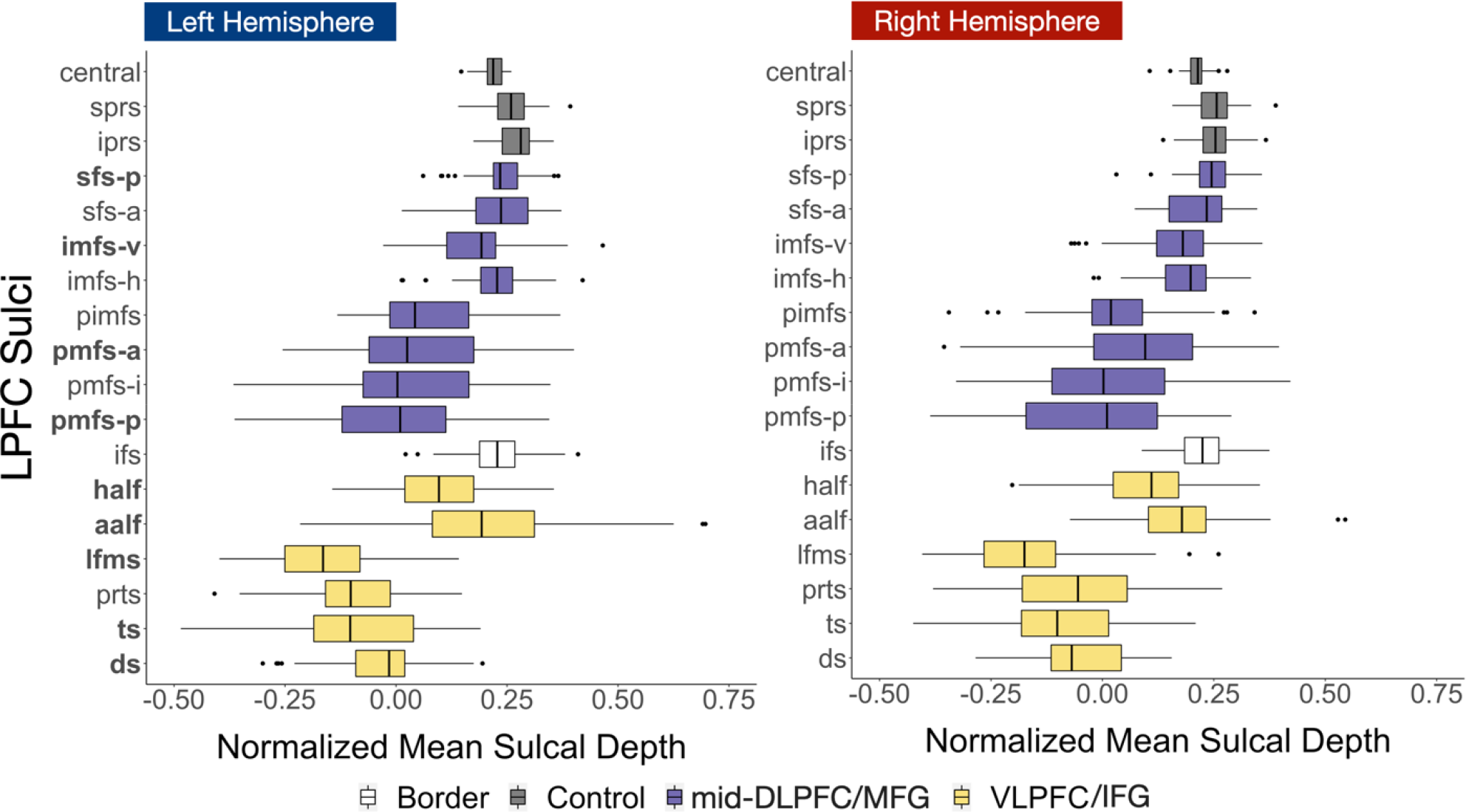
VLPFC/IFG sulci are shallower and more variable than MFG/mid-DLPFC sulci in a developmental cohort. Comparison of the normalized mean sulcal depths in the left and right hemispheres. Mean sulcal depth at each vertex in standard FreeSurfer units was normalized to the maximum depth within each hemisphere. Boxplots denoted in purple represent MFG/mid-DLPFC sulci, which are comparatively deeper and exhibit less variability in depth across participants. Boxplots in yellow represent IFG/VLPFC sulci, which are comparatively shallower and demonstrate more variability in sulcal depth across participants. Gray boxplots represent sulci not included in these verbal WM regions (cs, sprs, iprs) that are considered control sulci. The ifs (white) demarcates the boundary between the MFG/mid-DLPFC and IFG/VLPFC, and thus is not included in either verbal WM region. The sulci whose depths predict working memory manipulation – left hemisphere ds, ts, lfms, aalf, half, pmfs-p, pmfs-a, sfs-p, and imfs-v – are bolded on the y-axis.

#### 2.3.3. Characterization of Sulcal Morphology

For each individual, mean sulcal depth values were extracted for each sulcal label in each hemisphere by intersecting the label file with the .sulc file generated by FS using custom Python code (Miller et al., 2021; Voorhies et al, 2021). Raw depth metrics (standard FreeSurfer units) were computed in native space from the .sulc file generated in FreeSurfer 6.0.0. To account for differences in cortical depth across individuals and hemispheres, mean sulcal depth of each sulcus is reported as a proportion of maximum depth in each hemisphere (Voorhies et al., 2021). Mean cortical thickness of each sulcus was also considered as an additional metric to examine the extent to which the relationships between sulcal depth and behavior were specific or extended to other morphological features.

##### 2.3.3.1. Comparison between mid-DLPFC and VLPFC

To compare depth of sulci in the mid-DLPFC and VLPFC, we conducted a 2 way (verbal WM region x hemisphere) repeated measures analysis of variance (rm-ANOVA; Fig. 2). To assess variability between hemispheres and verbal WM region, we also conducted the same rm-ANOVA using standard deviation instead of mean sulcal depth. ANOVAs were computed in R with the *aov* function.

#### 2.3.4. Relating Sulcal Morphology and Behavior

##### 2.3.4.1. LASSO Regression and Cross-Validation

To assess the relationship between sulcal depth and working memory performance, we used a least absolute shrinkage and selection operator (LASSO) regression to select which of the 18 sulci, if any, are associated with Digit Span task scores. LASSO performs L1 regularization by applying a penalty, or shrinking parameter (α), to the absolute magnitude of the coefficients such that low coefficients are set to zero and eliminated from the model. In this way, LASSO can facilitate variable selection, leading to simplified models with increased interpretability and prediction accuracy (Heinze, 2018). We fit a LASSO regression separately for each hemisphere with each Digit Span task using a leave-one-subject-out cross-validation (LOOCV) procedure. In accordance with accepted methodological techniques, we tested a range of values for the shrinking parameter and (α) selected the value that minimized the error (LH: α = 0.01; Fig. 3) (Heinze et al., 2018).

**Figure 3.**
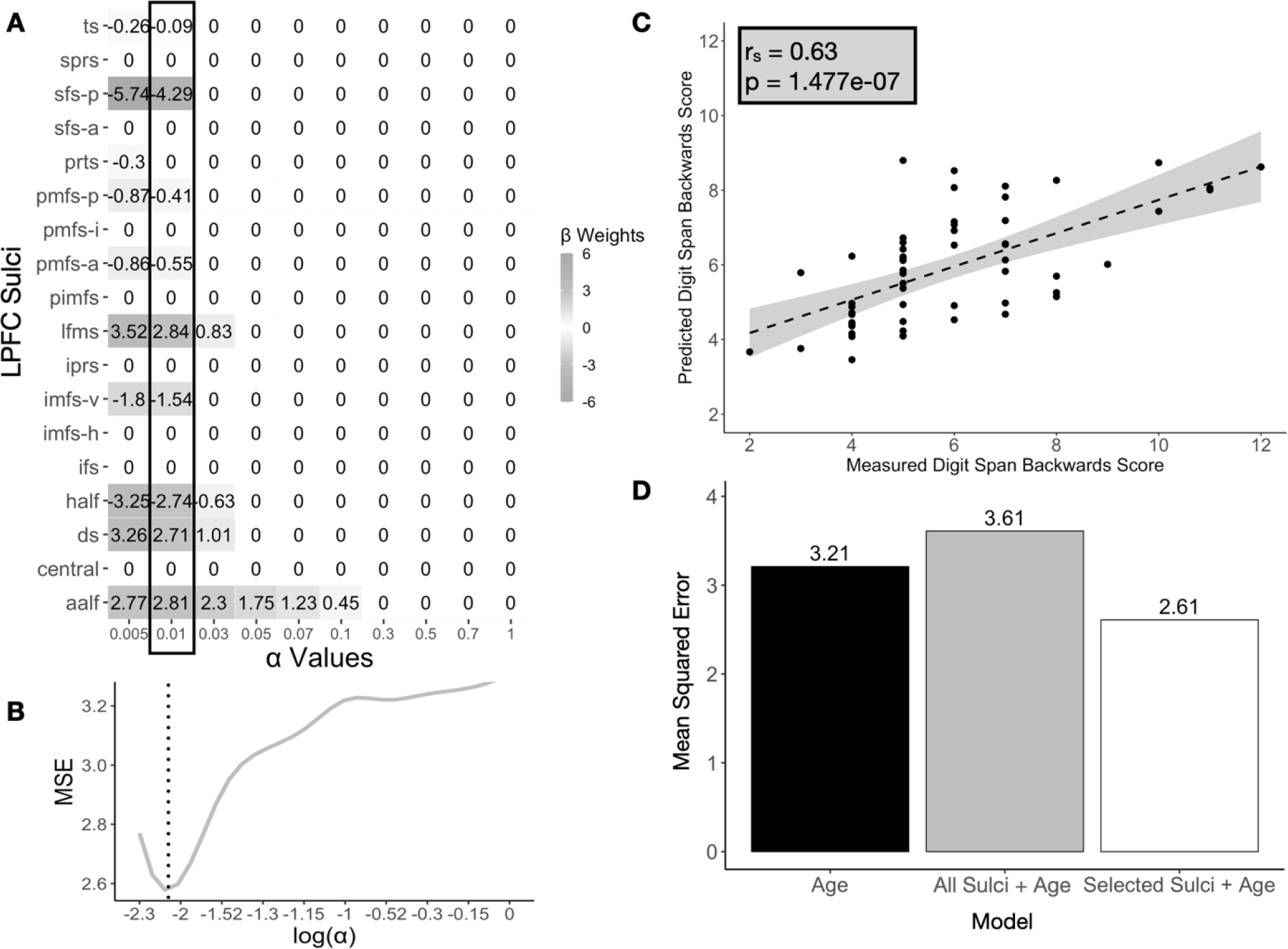
Performance and model fits relating sulcal depth to working memory performance. A. Beta-coefficients for each sulcus at a range of shrinking parameter (α) values resulting from the LASSO regression predicting Digits Backward score from sulcal depth in the left hemisphere. Highlighted box indicates coefficients at the chosen alpha-level. B. MSE at each alpha-level. We selected the value of α that minimized the Mean Squared Error (MSE; dotted line). C. Spearman’s Correlation (rs=0.63) between participants’ actual scores on Digits Backward and the predicted scores resulting from the LOOCV for the best performing model, which had only left hemisphere LASSO-selected sulci. D. Model comparison of the mean squared errors of a model with age as it only predictor (black), a model with all the left hemisphere sulci and age as predictors (gray), and a model with sulcal depth of selected sulci in the left hemisphere and age as predictors (white). The model with left hemisphere sulci selected by the LASSO regression (white) had the lowest mean squared error, performing the best.

##### 2.3.4.2. Model Comparisons

To specifically characterize the relationship between sulcal depth and verbal WM performance, we compared the models determined by the LASSO regression to two alternative nested models. Our alternative models only examined left hemisphere sulci as our initial models only identified sulci in the left hemisphere (Figure 3). As age is a factor in both sulcal development and verbal WM performance, we included age at the time of the assessment as an additional predictor in the models. Using the nine predictors of Digits Backward identified by the LASSO regression described above, we constructed a simplified linear model for the left hemisphere [1]. We refer to this simplified model as the *partial* model.

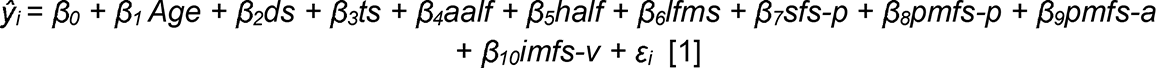

We compared the fit of the *partial* model with a *full* model that included all of the sulci within the left hemisphere, as well as age [2].

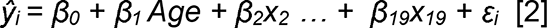

In the nested full model, x_2_ **-** x_19_ represent the sulcal depth of each of the 18 LPFC sulci in one hemisphere, and β_2_ **-** β_19_ represent the associated coefficients. To understand the role of age in any observed relationship, we also compared the full and partial models to a third nested model with age as the only predictor [3].

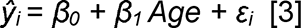

The model with the best fit was determined as having the lowest mean squared error (MSE) and the highest R-squared value. Linear models were fit using the SciKit-learn package in Python.

##### 2.3.4.3. Model Specificity

To ascertain whether or not any observed relationship between sulci and task performance generalized to other morphological features, we used mean cortical thickness instead of normalized mean sulcal depth as a predictor of Digit Span task scores. We then compared the fit of the best sulcal depth model to the cortical thickness model using the Akaike Information Criterion (AIC; Akaike, 1974). By comparing AIC scores, we could assess the relative performance of the two models. If the Δ greater than 2, it suggests an interpretable difference between models. If the Δ greater than 10, it suggests a strong difference between models, with the lower AIC value indicating the preferred model.

## 3. Results

For each participant, cortical surface reconstructions were generated using T1-weighted MPRAGE scans, and 18 LPFC sulci were manually labeled in each hemisphere. Depth and mean cortical thickness were calculated for each sulcus. To account for systematic individual and hemispheric differences in brain size, sulcal depth is calculated as a percentage of maximum depth in each hemisphere. This normalized sulcal depth is reported in standard Freesurfer units. To assess the relation between sulcal anatomy and verbal WM performance, we applied a data-driven approach (Voorhies et al., 2021). Below, we discuss the results of our approach in which we found that a) VLPFC sulci are shallower and more variable than mid-DLPFC sulci, b) sulcal depth is related to verbal WM manipulation, c) nine left-hemisphere LPFC sulci, but no right-hemisphere LPFC sulci defined in the present study predict manipulation scores, and d) these brain-behavior relations are not generalizable, as the cortical thickness of these nine sulci is not related to verbal WM performance.

### 3.1. Tertiary sulci are consistently identifiable in LPFC of 6-18 year-olds and VLPFC sulci are shallower and more variable than mid-DLPFC sulci

17 of the 18 LPFC sulci were identifiable in both hemispheres in all participants. Only the *pimfs*, a tertiary sulcus, was unidentifiable in the right hemisphere for two participants and in the left hemisphere for one participant (Supplementary Fig. 1). A two-way repeated measures analysis of variance (rm-ANOVA) was conducted to statistically test for differences between verbal WM region (MFG/mid-DLPFC, IFG/VLPFC) and hemisphere. The rm-ANOVA revealed a main effect of verbal WM region (F(1,56) = 295.7, *p<*0.001), showing that VLPFC were more shallow than mid-DLPFC sulci (*M*_VLPFC_= -0.01; *M*_mid-DLPFC_ = 0.12; Fig. 2). However, there was no effect of hemisphere on sulcal depth (F(1,56) = 1.84, *p =* 0.18) or a hemisphere x verbal WM region interaction (F(1,56) = 0.26, *p =* 0.61). To explore the variability between verbal WM regions, we repeated this rm-ANOVA substituting mean sulcal depth with standard deviation of sulcal depth. This analysis revealed a main effect of verbal WM region (F(1,56) = 6.18, *p=*0.01), showing that the depths of VLPFC sulci were more variable than the depths of mid-DLPFC sulci (*SD*_VLPFC_ = 0.18; *SD*_mid-DLFPC_ = 0.16). We also observed an effect of hemisphere on standard deviation of sulcal depth (F(1,56) = 8.44, *p <* 0.01), but did not observe a hemisphere x verbal WM region interaction (F(1,56) = 0.073, *p =* 0.79).

### 3.2. Sulcal depth is associated with working memory manipulation, but not maintenance

To characterize the relationship between sulcal depth and the two aspects of working memory (manipulation and maintenance), we applied feature selection to determine which sulci, if any, were associated with either maintenance (Digits Forward) or manipulation (Digits Backward). To do so, we implemented a LASSO regression relating the depths of 18 sulci (Fig. 1) as well as age, to scores on the two tasks (Materials and Methods). A LASSO regression not only allows us to select sulci in a data-driven manner, but also improves the generalizability of a model and prevents overfitting, particularly in cases where there are 10 &#x003C; x &#x003E; 25 predictors (Heinze et al., 2018).

We assessed the relationship between sulcal depth and maintenance and manipulation separately in each hemisphere. Age was included as an additional predictor in all models. This approach revealed a significant association between normalized mean sulcal depth and Digits Backward score in the left (*R^2^* = 0.40, *MSE* =2.61, α = 0.01), but not the right (*R^2^* = 0.25, *MSE* = 3.22, α = 0.1), hemisphere. Neither left nor right hemisphere sulcal depth was related to Digits Forward score (LH: *R^2^* = 0.26, *MSE* = 3.27, α = 0.07; RH: *R^2^* = 0.26, *MSE* = 3.27, α = 0.07), even though Digits Forward and Backward scores were correlated across participants (*r* = 0.51, *p* < 0.001). As predicted, age was also associated with working memory performance for both the Digits Backward condition (*R^2^* = 0.25, *MSE* = 3.21) and Digits Forward condition (*R^2^* = 0.26, *MSE* = 3.21). There was a positive relationship between task performance and age: the older the participant, the better the performance on Digits Backward (β_age_= 0.34) and Digits Forward (β _age_= 0.35) tasks.

### 3.3. The depths of nine left hemisphere LPFC sulci predict working memory manipulation task performance

Examining the coefficients of the manipulation models revealed that nine out of 18 left hemisphere LPFC sulci, but none of the right hemisphere sulci, were related to performance for the Digits Backward task. In left IFG/VLPFC, the *ds, ts, lfms, aalf,* and *half* were predictors in the model (Fig. 3a, Fig. 4). In left MFG/mid-DLPFC, the *sfs-p, imfs-v, pmfs-p,* and *pmfs-a* were additionally predictive (Fig. 3a, Fig. 4). These sulci were selected based on the results of the LASSO regressions. Six of the nine sulci exhibited negative relationships with verbal WM manipulation performance, whereas three others demonstrated positive relationships with verbal WM manipulation performance (Fig. 3).

**Figure 4.**
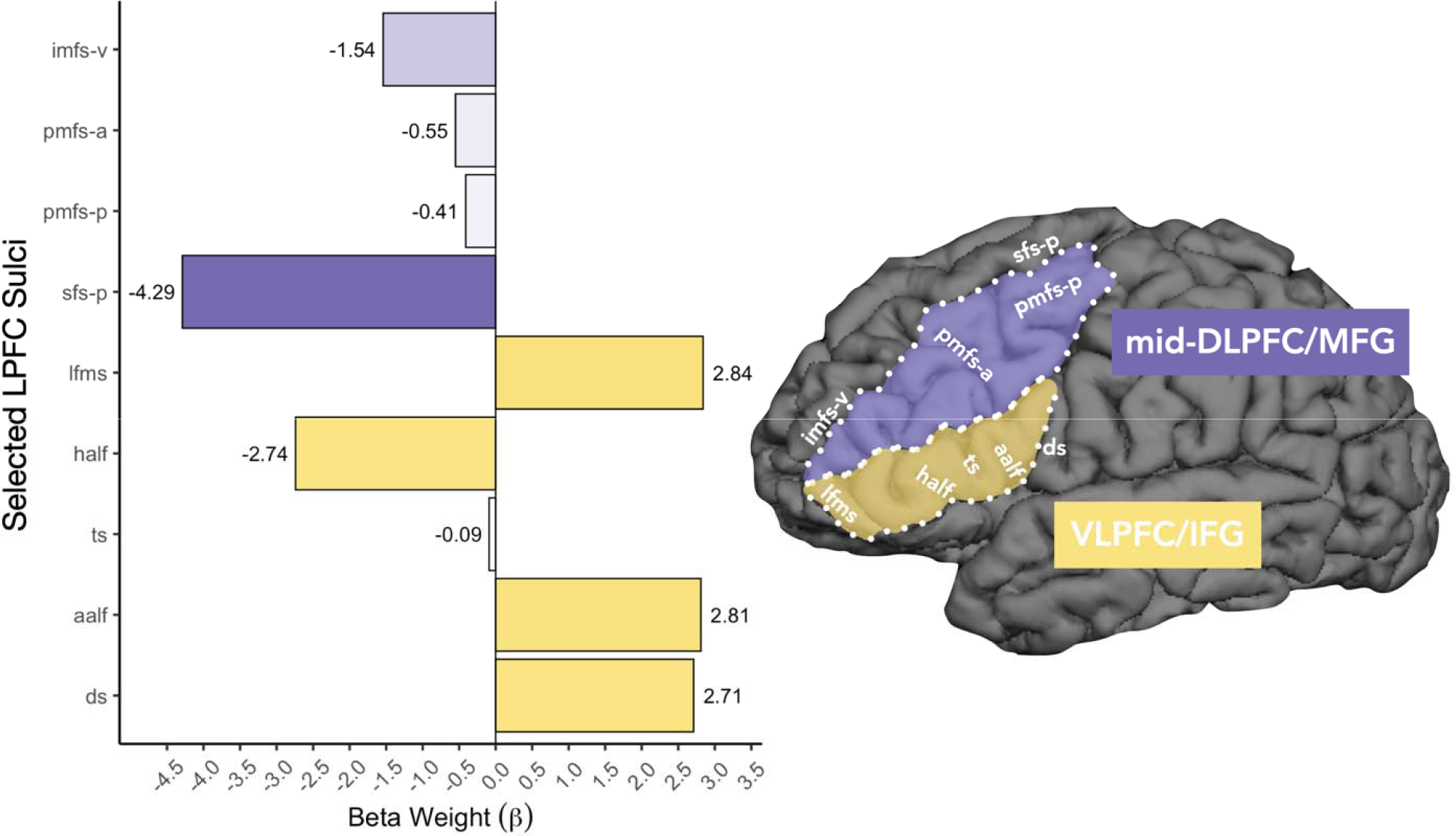
The relationship among function, sulcal morphology, and working memory skills in the developing lateral prefrontal cortex. Left: Barplot comparing the beta weights for Digits Backward for each of the nine left hemisphere sulci identified by the model. Negative and positive beta coefficients represent negative and positive relationships between sulcal depth and tas performance, respectively. All four MFG/mid-DLPFC (purple) sulci show negative relationships with verbal WM manipulation, while 3/5 IFG/VLPFC (yellow) sulci show positive relationships with verbal WM manipulation. Darker shades represent stronger beta coefficients. These values are also shown in Figure 3A. Right: An example cortical surface reconstruction of a left hemisphere in an example 15-year-old (n039t2). Sulcal depth predicted working memory skills in the ds, ts, aalf, half, lfms, pmfs-p, pmfs-a, sfs-p, and imfs-v (white labels). The ds, aalf, half, lfms, imfs-v, and sfs-p form the boundaries of the functional clusters preferentially coding the serial order of visual stimuli in working memory identified by Amiez and Petrides (2007) and have stronger relationships with verbal WM manipulation than the ts, pmfs-a, and pmfs-p, which fall within these functional borders. Purple: MFG/mid-DLPFC based on macroanatomical definition of mid-DLPFC in Amiez and Petrides (2007). Yellow: IFG/VLPFC based on macroanatomical definitio of VLPFC in Amiez and Petrides (2007).

To further examine the relationship between the depth of these specific sulci and verbal WM manipulation, we used the partial model derived from our LASSO regression with LOOCV to predict performance. The partial model only used the mean sulcal depths of left hemisphere *ds, ts, lfms, aalf, half, sfs-p, imfs-v, pmfs-p,* and *pmfs-a* as predictors of Digits Backward scores in the LOOCV linear regression. Age was also included as a predictor in the model. The results of the LOOCV linear regression confirmed that the partial model with these nine sulci significantly predicted Digits Backward scores (*r_s_* = 0.63, *p <* 0.0000001; Fig. 3C). When compared to an alternative nested model which included all the LPFC sulci in the left hemisphere, we found that the addition of the other nine sulci did not improve the fit of the model (*R^2^* = 0.16, *MSE* = 3.61; Fig. 3D). This comparison was consistent with the predictions of the LASSO regression.

To ascertain that the relationship between sulcal depth and verbal WM manipulation performance was not driven by age (see Supplementary Fig. 3 for relationship between age and depth of each sulcus), we additionally compared the left hemisphere partial model to another alternative nested model with age as the sole predictor (*R^2^* =0.26, *MSE*=3.21; Fig. 3D). Age at the time of assessment was positively correlated with scores on the Digits Backward task (*r* = 0.57, p < 0.001). However, when compared to the age model, the left hemisphere partial model (*MSE* = 2.61) showed a lower mean squared error. Thus, age alone did not explain the results of the partial model. The inclusion of the selected sulci improved prediction of Digits Backward scores above and beyond age.

### 3.4. Sulcal depth, not sulcal cortical thickness, is associated with verbal WM performance

To assess whether the association between sulcal depth and verbal WM manipulation performance extended to other morphological features of the sulci included in the partial models, we also used a LASSO approach with mean cortical thickness of each sulcus as the predictor. Again, age was included as an additional predictor in the left hemisphere partial models. We found a slight positive relationship between mean sulcal cortical thickness of the nine left hemisphere sulci and Digits Backward scores (*R^2^* = 0.04, *MSE* = 4.12). Nevertheless, the sulcal cortical thickness model did not perform better than age in predicting Digits Backward scores.

Additionally, sulcal depth was a better predictor of verbal WM manipulation than was sulcal cortical thickness, as assessed by the Akaike Information Criterion AIC = 27.48; Akaike, 1974). If the Δ is greater than 2, it suggests an interpretable difference between models; the lower AIC value indicates the preferred model (Wagenmakers and Farrell, 2004; Burnham and Anderson, 2004).

## 4. Discussion

To our knowledge, this is the first study to examine the relationship between mid-DLPFC and VLPFC tertiary sulcal morphology and working memory (verbal WM) performance in a developmental cohort. Implementing a data-driven approach with cross-validation, we showed that there is a relationship between sulcal depth and verbal WM performance: the depth of nine left hemisphere sulci predicted verbal WM maintenance and manipulation on the challenging Digits Span Backwards task. By contrast, none of the sulci predicted pure WM maintenance, as measured by the Digits Span Forwards task.

As noted previously, our study addressed three main questions. First, we sought to test for a relationship between verbal WM and mean depth of LPFC sulci. Our results support this hypothesis, showing an anatomical-behavioral relationship for numerous LPFC sulci. Second, we asked whether any such relationships differed as a function of sulcal type, hemisphere, and/or task demands. We found relationships for both shallow/tertiary and deep/non-tertiary sulci. However, there was a hemispheric effect: all of the sulci implicated in verbal WM were localized in the left, language-dominant, hemisphere. We also found an effect of task demands: we only observed relationships between sulcal depth and performance when the task required manipulation of items in WM. Third, we asked whether we could construct a model to predict an individual’s verbal WM task score from sulcal depth. Indeed, using a LASSO regression to select the sulci most related to performance and a LOOCV linear regression to predict scores on the task, we were able to construct a model including age and the nine left LPFC sulci that better predicted scores compared to separate models with either all LPFC sulci or age alone.

The sulci predicting verbal WM manipulation included six out of ten of the tertiary sulci labeled in this study, as well as three of the eight primary sulci. While greater sulcal depth predicted better performance for three sulci, shallower sulcal depth predicted better performance for six sulci. In the sections below, we discuss (i) how these findings contribute to building empirical support for a classic hypothesis proposing that features of tertiary sulci in association cortices would be functionally and behaviorally relevant for cognition, (ii) potential underlying mechanisms contributing to the relationship between the depths of tertiary sulci and human behavior, and (iii) future directions and limitations of our study.

### 4.1. Sulcal depth in left hemisphere LPFC predicts verbal WM manipulation: Linking structure, function, cognition, and lateralization with theoretical insights

Of the nine left hemisphere sulci whose depths predicted verbal WM performance, a majority were tertiary sulci, consistent with Sanides’ theory (1962, 1964). Specifically, mirroring the fact that primary sulci, which emerge early in gestation, serve as landmarks in primary sensory cortices, Sanides posited that tertiary sulci, which emerge late in gestation, serve as landmarks in association cortices, which also show a protracted development. He further proposed that the late emergence and continued postnatal morphological development of tertiary sulci is likely related to cognitive skills associated with LPFC that also show a protracted development. While our results support this hypothesis by showing that tertiary sulci are behaviorally meaningful, we did not find that *only* tertiary sulci were linked to cognitive performance. This fits with previous findings that also showed relationships between non-tertiary sulcal morphology in other cortical locations and cognitive performance in children (Roell et al., 2021). Interestingly, the locations of the nine sulci in the left hemisphere identified by our model align with the functional definitions of mid-DLPFC and mid-VLPFC as identified by Amiez and Petrides (2007; Fig. 4 - right). That is, the *ds, aalf, half, lfms, imfs-v*, and *sfs-p* form the boundaries of the functional clusters preferentially coding the serial order of visual stimuli in working memory identified by Amiez and Petrides (2007). These sulci also have stronger relationships with verbal WM manipulation than the ts, pmfs-a, and pmfs-p, which fall within these functional borders. This link between present and prior findings suggests that a subset of sulci identified by our model-based approach are potential functional landmarks, which can be examined in future studies.

The laterality effects identified in the present study are consistent with extensive neuropsychological and neuroimaging evidence that verbal WM tasks rely heavily on the left hemisphere (e.g., Black, 1986a, 1986b; Laures-Gore et al., 2011; Paulesu et al., 1993; Smith and Jonides, 1998). With regards to LPFC and the same tasks used in the present study, Barbey and colleagues (2013) showed that left, but not right, DLPFC lesions resulted in deficits in verbal WM manipulation; neither left nor right DLPFC lesions affected verbal WM maintenance. Together, these findings suggest that individual differences in sulcal development impact the functional organization of LPFC-dependent cognitive function identified here and in previous work (Garrison et al., 2015; Im et al., 2008; Voorhies et al., 2021).

### 4.2. Underlying anatomical mechanism(s) likely contributing to the relationship between sulcal depth and human behavior: Short-range connections and local gyrification?

We have recently proposed that deeper tertiary sulci in LPFC could be indicative of short-range connections that function to pull regions closer together and, in turn, are associated with greater efficiency of neural processing by decreasing the distance between LPFC regions (Voorhies et al., 2021). This heightened neural efficiency could manifest as improved behavioral performance. This mechanistic hypothesis was proposed based on recent and classic findings showing that a) short white-matter fibers extend from the deepest points of sulci into the white matter (Reveley et al., 2015) and b) there is a relationship between tertiary sulci in LPFC and myelination (Miller et al., 2021a,b; Sanides, 1962). The present findings build on this proposal by showing that a combination of shallower and deeper sulci predicts verbal WM performance.

Because in some cases we found a negative rather than positive relation between sulcal depth and cognitive performance, we speculate that additional anatomical mechanisms, such as those related to the development of neighboring sulci or white matter tracts, are likely at play. For instance, several researchers (Armstrong et al., 1995; Connolly, 1950; Zilles et al., 2013) qualitatively noted that the sizes and depths of sulci seemingly counterbalance those of nearby sulcal neighbors. Thus, a shallow, short sulcus would compensate for a particularly long and deep nearby sulcus, rendering the overall degree of cortical folding within a given region approximately equal (Armstrong et al., 1995; Connolly, 1950; Zilles et al., 2013). Given this hypothesis, we propose that a relatively deeper tertiary sulcus, which may have stronger short-range white matter connections as proposed previously (Voorhies et al., 2021), may be close to sulci that are relatively shallower, thereby preserving the degree of local cortical folding. This proposal builds on a recent modification of the compensation theory of cortical folding that proposes to also incorporate local morphological features (Natu et al., 2020). Altogether, our findings begin to build a multimodal mechanistic neuroanatomical understanding underlying the complex relationship between sulcal depth and cognition relative to other anatomical features, which importantly makes explicit, testable predictions for future studies.

### 4.3. Future directions and limitations

The main limitation of our study is the labor-intensive process of manually identifying sulci, which limits the number of participants. Current methods implement deep learning algorithms to automatically identify primary and secondary sulci in LPFC (Hao et al., 2020). Thus, modifications to these algorithms to include tertiary sulci would make it possible to expand sample sizes in future studies examining the relationship between the morphology of tertiary sulci and cognition. Ongoing work is already underway to develop deep learning algorithms to accurately define tertiary sulci automatically in individual participants, and initial results are promising (Borne et al., 2020; Lyu et al., 2021).

Additionally, verbal WM depends on a distributed neural system; thus, it is likely that verbal WM performance is also related to sulcal variation elsewhere in the brain, such as lateral parietal cortex (Goldman-Rakic, 1990; Klingberg, 2006; Smith and Jonides, 1998; Tamnes et al., 2013). Because maintenance matures more quickly than manipulation (Gathercole et al., 2004), linked to the differential developmental trajectory of the IFG and MFG (Crone et al., 2006), examining the relationship between sulcal depth and verbal WM at an earlier age might also reveal the involvement of other sulci in maintenance, and in verbal WM more generally. Furthermore, although it is common to relate brain structure to performance on individual tasks, as we have done here, the use of a latent construct or composite score derived from multiple verbal WM measures would permit us to further assess the generalizability of our cross-validated results (see Bollen, 2002).

To explore why various sulci showed opposite relationships between depth and performance, future work should also examine a) relationships between the development of neighboring sulci and b) the underlying white matter tracts linking these sulci. The former will allow us to understand if compensation, whereby shallower or shorter sulci balance nearby longer or deeper sulci, is at play. The latter will reveal if sulcal depth is linked to white matter connections and increased neural efficiency, which may be related to individual differences in cognitive performance.

Finally, future studies involving large sample sizes should also consider how other variables like demographics, genetics, and early environment also contribute to this relationship between sulcal depth and cognition. Semi-automation of tertiary sulcal definitions further render the feasibility of larger-scale investigations (Borne et al., 2020; Lyu et al., 2021).

### 4.4. Conclusion

These findings highlight the behavioral significance of individual variability in sulcal morphology. Some of these sulci may serve as boundaries for functional regions engaged in verbal WM manipulation. These results begin to shed light on the complex relationship among sulcal morphology in LPFC, functional parcellations of LPFC, and verbal WM skills in children and adolescents. More broadly, these and our prior findings based on a largely overlapping pediatric MRI dataset (Voorhies et al., 2021) contribute to an increasing body of work in adults that also empirically support Sanides’ hypothesis that tertiary sulci serve as functional and cognitive landmarks in association cortices. Taken as a whole, this emerging body of research indicates the importance of studying tertiary sulci to better understand the relationships among brain structure, brain function, and behavior.

## Acknowledgments

We thank Ishana Raghuram, Chahat Mittal, and Anmol Gill for assistance with sulcal labeling and former members of the Bunge laboratory for assistance with data collection, and the families who participated in the study.

## Supplementary Materials

**Supplementary Figure 1.**
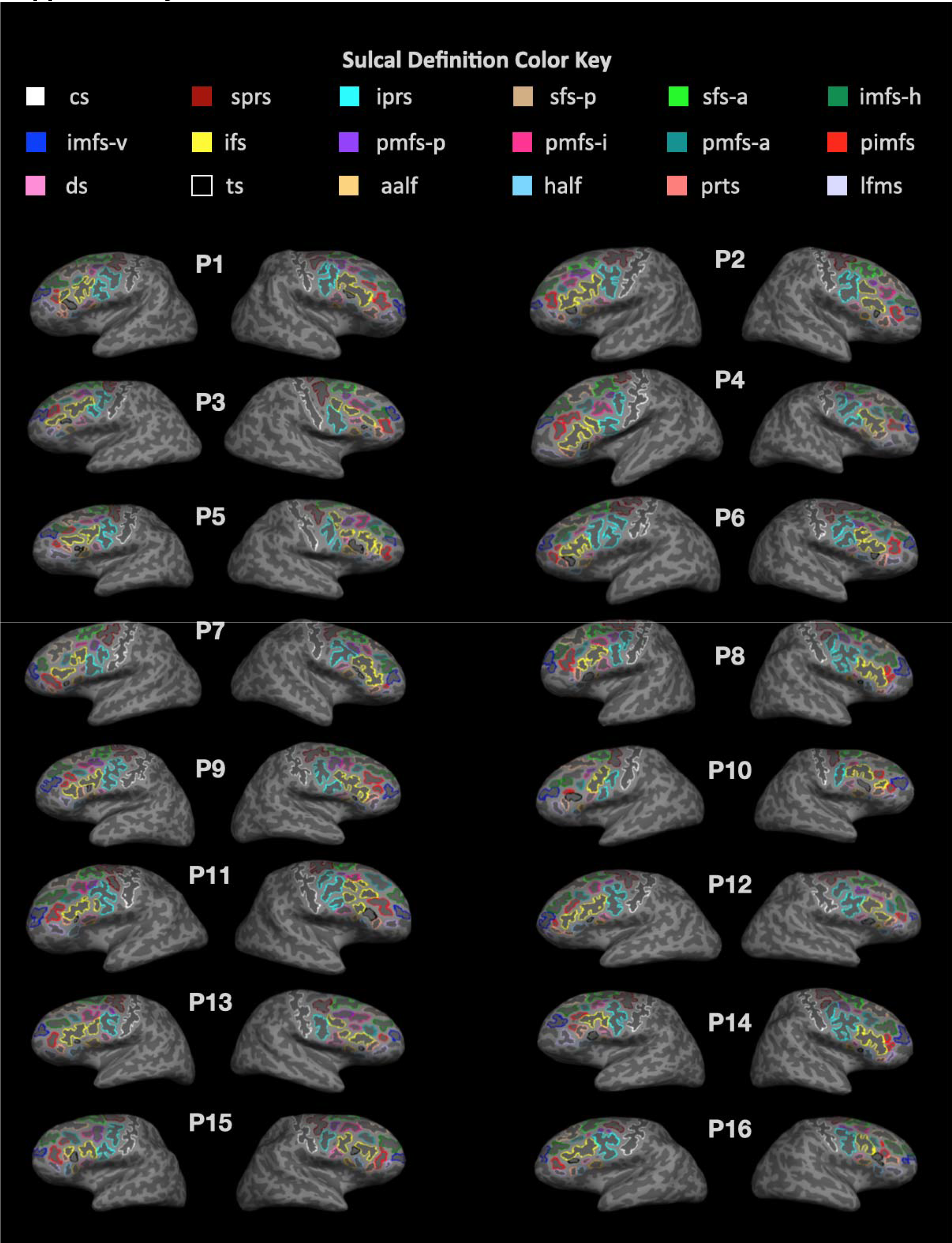
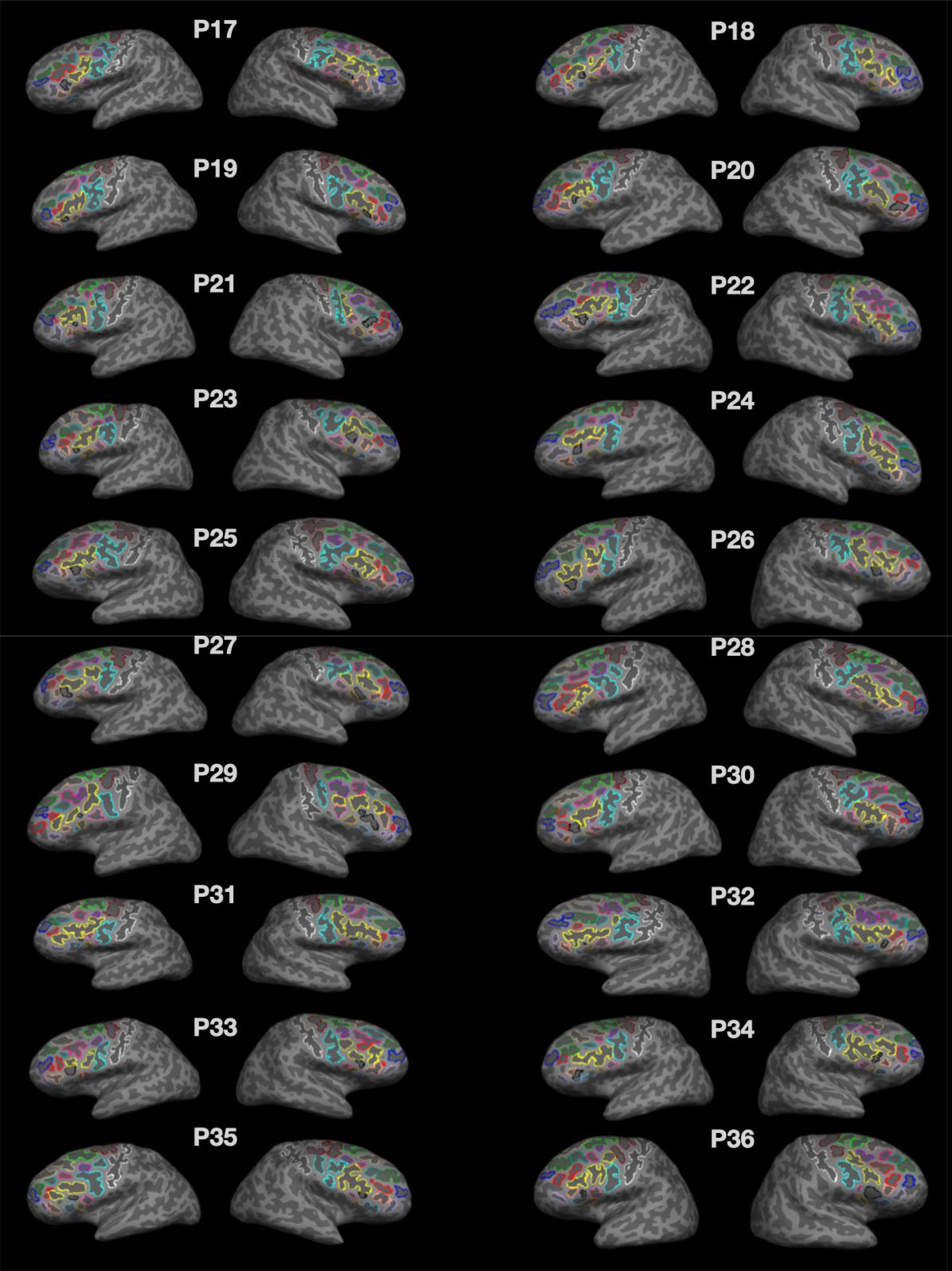
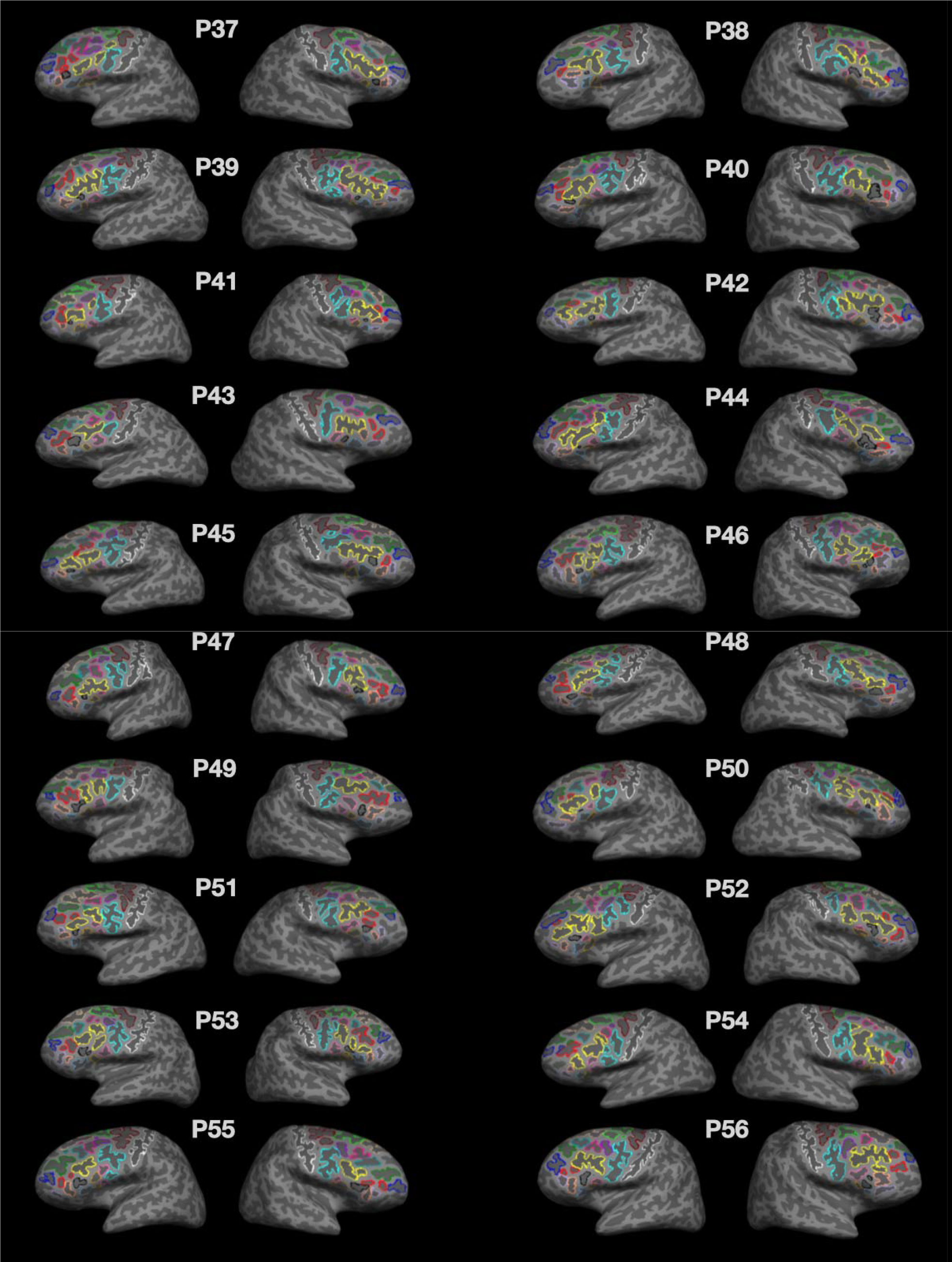
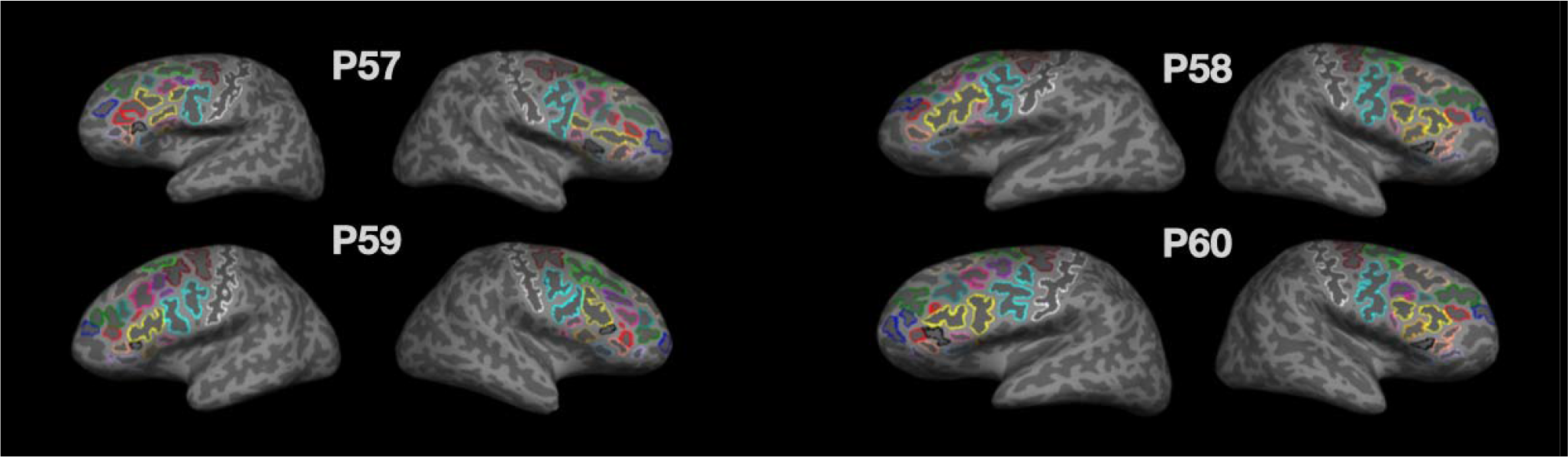
Manual sulcal labels in right and left hemispheres for each participant. Manually labeled sulci on inflated cortical surface reconstructions for left and right hemispheres of each participant (N=60). 18 individual sulci are outlined by color indicated by the key. The *pimfs* can include 1 or 2 components, or not be present. For participants with two *pimfs* components, the label is composed of both components.

**Supplementary Figure 2.**
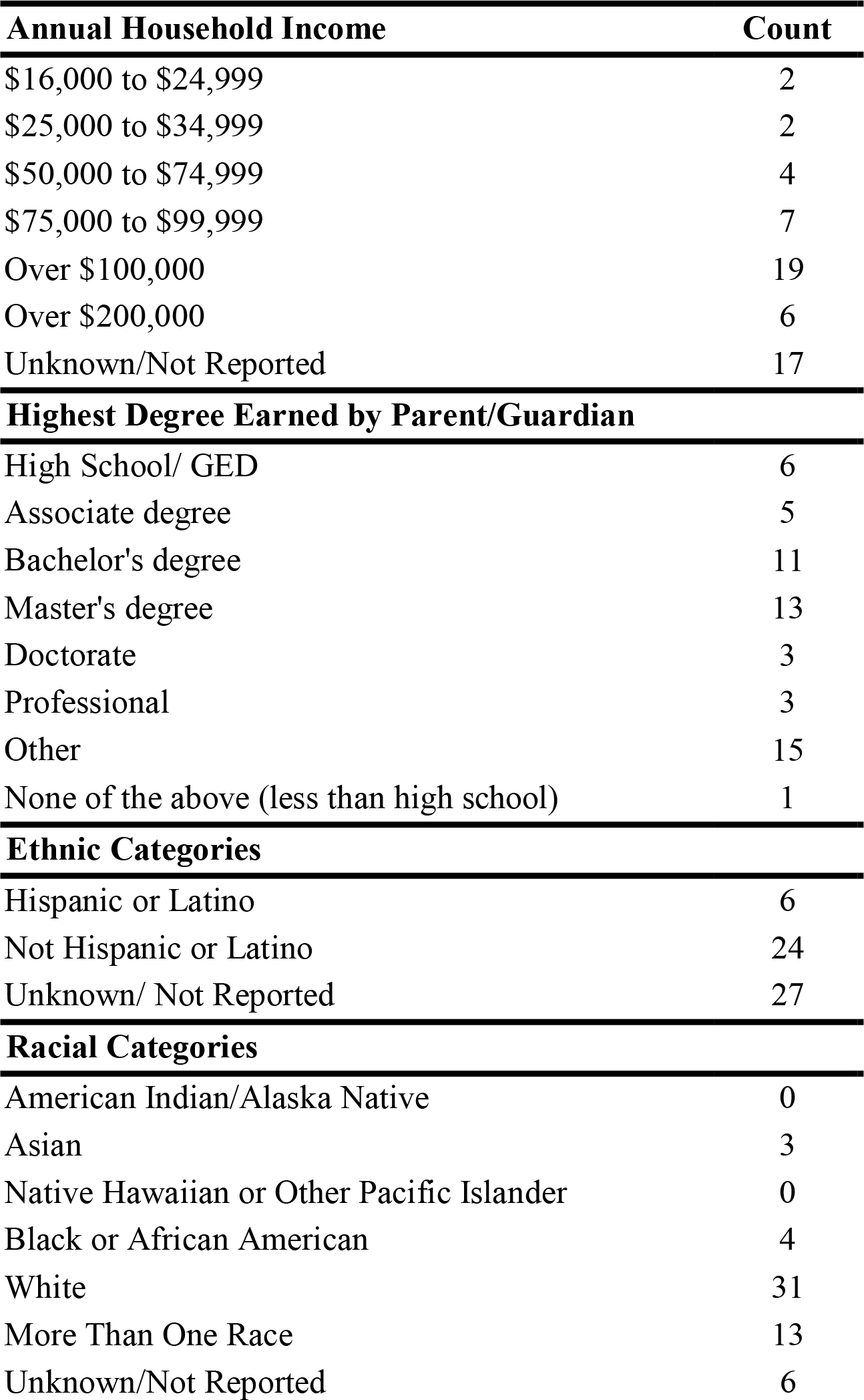
Summary of participant demographics. Parent/Guardian reported Family income, Education, Race, and Ethnicity are summarized for all participants included in the behavioral analyses (N=57).

**Supplementary Figure 3.**
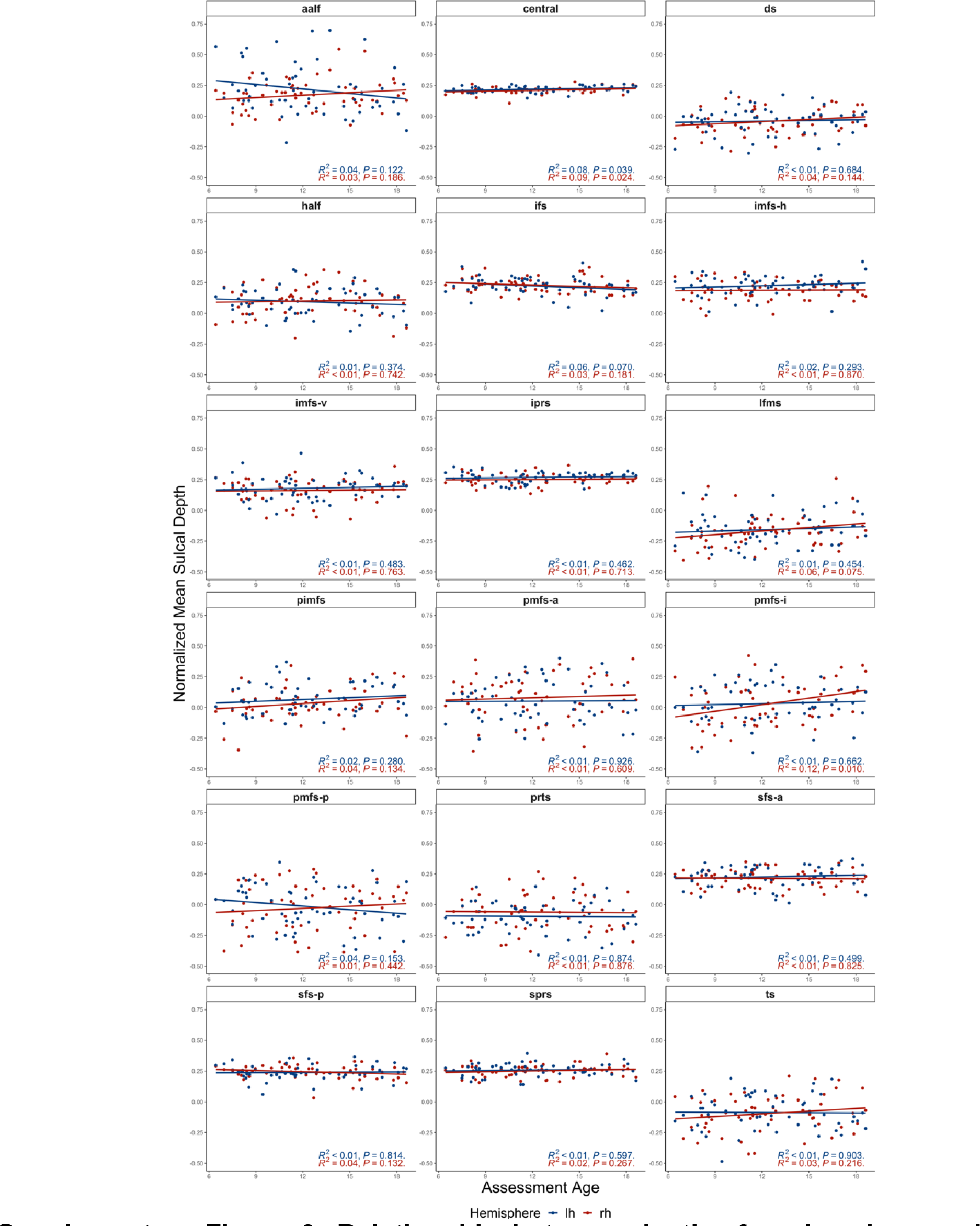
Relationship between depth of each sulcus and age. Relationship of the normalized mean depth of each of the 18 sulci in the left (blue) and right (red) hemispheres with age. R-squared values and p-values are reported at the bottom right of each subplot.

